# Can early assessment of hand grip strength in elderly hip fracture patients predict functional outcome?

**DOI:** 10.1101/557371

**Authors:** Ivan Selakovic, Emilija Dubljanin-Raspopovic, Ljiljana Markovic-Denic, Vuk Marusic, Andja Cirkovic, Marko Kadija, Sanja Tomanovic-Vujadinovic, Goran Tulic

## Abstract

Decreased muscle strength is not only a risk factor for hip fracture in elderly patients, but plays a role in recovery of physical function. Our aim was to assess the role of grip strength measured early after hip fracture, and classified according to the EWGSOP2 criteria in predicting short- and long-term functional recovery. One hundred ninety-one patients with acute hip fracture consecutively admitted to an orthopaedic hospital have been selected. A multidimensional geriatric assessment evaluating sociodemographic variables, cognitive status, functional status and quality of life prior to fracture, as well as perioperative variables were performed. Follow-ups at 3 and 6 months after surgery were carried out to evaluate functional recovery. Multivariate regression models were used to assess the predictive role of handgrip strength. The mean age of the participants was 80.3 ±6.8 years. Thirty-five percent of our patients with clinically relevant hand grip strength weakness were significantly older, more often female, had a lower BMI, and were of worse physical health. They also had a lower cognitive level, lower Barthel index, and lower EQ5D scores before fracture. Multivariate regression analysis adjusted for age and gender revealed that hand grip weakness was an independent predictor of worse functional outcome at 3 and 6 months after hip fracture for both genders and in all age populations. Our study supports the prognostic role of hand grip strength assessed at hospital admission in patients with hip fracture. Thus, clinicians should be encouraged to include hand grip assessment in their evaluation of hip fracture patients in the acute setting in order to optimize treatment of high-risk individuals.

## Introduction

Sustaining a hip fracture is considered one of the most fatal fractures for elderly people that leads to impaired function, and increased morbidity and mortality, and high financial liability. These facts challenge clinicians in identifying patients at risk of worse outcome early in the course of hip fracture treatment, in order to set realistic rehabilitation goals, optimize perioperative care, and define optimal rehabilitation strategies in order to reduce devastating outcomes.

Functional evaluation in patients with hip fracture is an essential part of multidimensional assessment, and has an important prognostic value. Muscle weakness is considered a key element of frailty [1] and, increasingly, of sarcopenia [2, 3]. It is believed that sarcopenia not only enhances fracture risk, but also increases the risk of poor functional outcome after hip fracture [4]. Reduced muscle strength makes it more difficult to regain lost balance and decreases the mechanical loading of the skeleton leading to reduced adaptive bone remodeling [5, 6]. Hand grip strength (HGS) assessment is an objective measure of overall body muscle strength and physical function [7, 8], an important measure for frailty [9], and sarcopenia [10, 11]. Various studies have shown the prognostic value of hand grip strength in patients with hip fracture [12–17]. However, very few have been carried out in the acute phase [16, 17], but there are no studies which have been carried out using the European Working Group on Sarcopenia in Older People 2 (EWGSOP2) criteria [3] to define clinically relevant hand grip weakness.

The aim of our study was to assess the EWGSOP2 threshold for grip strength assessed at admission to hospital after hip fracture to predict short- and long-term functional recovery. We hypothesized that levels of grip strength below the EWGSOP 2 thresholds measured in the first 48h after hip fracture could predict an unfavorable short- and long-term functional outcome.

## Materials and methods

### Study design

All adult patients 65 years or older with an acute hip fracture who were admitted consecutively to an university associated orthopedic hospital in Serbia between March 1st 2017 and February 28th 2018 were enrolled in an open, prospective, observational cohort study. All patients with pathologic fractures, major concomitant injuries, multiple trauma, malignant diseases, imminent death as a result of an end-stage disease, inability to walk before fracture, and nonoperative treatment resulting from high surgical risk were excluded. Furthermore, patients with severe cognitive impairment, as well as patients with hand weakness as a consequence of previous neurologic disorders or hand injuries also were excluded. During the study period, 551 patients had hip fractures and were examined for eligibility. One hundred ninety-one patients were confirmed eligible and were included in the study. All patients gave written informed consent to participate in the study. The study was conducted according to the Helsinki Declaration and approved by the University’s institutional review board.

## Measures

### Baseline evaluation

We assessed all subjects through standardized patient interview with respect to sociodemographic variables (age, sex, marital status, preinjury living conditions), cognitive level, handgrip strength, prefracture functional level, and health related quality of life within 24h of admission. We also recorded perioperative variables during the primary hospital stay, such as comorbidity level, waiting time for surgery, type of fracture, surgical method, type of anesthesia, and presence of postoperative complications, and length of stay (LOS).

Cognitive level was assessed with the Short Portable Mental Status Questionnaire (SPMSQ) [18]. The 10-item questionnaire classifies the patient’s cognitive level depending on the number of correct answers as lucid (8–10), mild to moderate cognitive dysfunction (3–7), and severe cognitive dysfunction (0–2). Handgrip strength was measured using a JAMAR hand dynamometer (Model BK-7498, Fed Sammons Inc, Brookfield, III). Patients were in the supine position, and encouraged to exhibit the greatest possible force [19]. The best recorded of 3 attempts of maximal voluntary contraction performed at 1-minute intervals of the dominant hand was considered for analysis. Hand grip strength measurements less than 16 kg in women and 27 kg in men were considered cut-points for the diagnosis of sarcopenia according to the revised EWGSOP2 criteria [3]. The pre-fracture functional status 2 weeks before hospital admission was assessed by the Barthel index [20]. The Barthel index measures performance in basic activities of daily living; its score ranges from 0 (total dependence) to 100 (total independence) [21]. General health related quality of life was measured with the EQ5D scale, which consists of a five-level response for five domains related to daily activities, mobility, self-care, usual activities, pain and discomfort, anxiety and depression [22]. Responses to the health status classification system are converted into an overall score using a published utility algorithm for the UK population [23].

We used the Charlson comorbidity index (CCI) to categorize comorbidities [24]. Patients were divided into three groups: without and mild, with CCI scores of 1–2; moderate, with CCI scores of 3–4; and severe, with CCI scores ≥5.

All patients with femoral neck fractures (84 patients (43.9%)) underwent bipolar hemiarthroplasty, whereas all patients with intertrochanteric (92 patients (48,2%)) and subtrochanteric fractures (15 patients (7.9%)) underwent open reduction and internal fixation (ORIF). In all patient early assisted ambulation was encouraged on the first postoperative day with weightbearing as tolerated, and all patients followed a standardized postoperative rehabilitation program.

### Outcomes

Functional status after 3 and 6 months was evaluated using the Barthel index score. The information was collected by phone interview. Data from patients who died or were lost before the first and second follow-up respectively were excluded from the study. For the analysis of Barthel index 3 months postoperatively the sample size included 160 patient (22 (11.5%) died; 9 (4.7%) were lost to follow-up). Analysis of outcomes six months after the fracture was performed on 154 patients (27 (14.1%) died, 10 (5.3%) were lost to follow-up).

### Statistical analysis

Continuous variables are presented in terms of mean values with SD or median and interquartile range depending on Kolmogorov-Smirnov test of distribution normality. Categorical values are summarized as absolute frequencies and percentages. To compare patients with two different categories of grip strength a *t* test was performed for the continuous variables and a Mann-Whitney U test for ordinal variables.

In order to detect potential and independent predictors of recovery expressed as Barthel index scores after 3 and 6 months, univariate and then multivariate linear regression with collinearity diagnostic (VIF method used; variables with VIF > 4 were excluded from multivariate models) was used. Both multivariate models were adjusted for age and gender.

The significance level for all statistical tests was set at 0.05. All analyses were performed using the SPSS Inc. Released 2008. SPSS Statistics for Windows, Version 17.0. Chicago: SPSS Inc.

## Results

Our cohort consisted of 191 patients aged 66 to 97 years. The mean age was 80.3 ±6.8 years, and 77.0% of our cohort were women. The mean HGS in our cohort was 20.5 ±6.8 kg. Sixty-six (34.6%) patients had clinically relevant hand grip weakness. Those patients were significantly older, more often female, had a lower BMI, and were of worse physical health. They also had a lower cognitive level, lower Barthel index scores and lower EQ5D scores before fracture. Patients with weaker grip strength were more often operated in general anesthesia (Table 1).

**TABLE 1.** Socio-demographic and baseline pre- and perioperative characteristics of the participants

|  | **Women With HGS < 16 kg**<br>**Men With HGS < 27 kg**<br>**N=66 (34.6%)** | **Women With HGS ≥ 16 kg**<br>**Men With HGS ≥ 27kg**<br>**N=125 (65.4%)** | **p** |
| --- | --- | --- | --- |
| **Age (year)\*** | 83.53 ± 6.16 | 78.52 ± 6.50 | <0.001 |
| **Gender\*\*** |  |  |  |
| Male | 21 (31.8%) | 23 (18.4%) | 0.036 |
| Female | 45 (68.2%) | 102 (81.6%) |  |
| **Marital status\*\*** |  |  |  |
| Other | 40 (61.5%) | 76 (61.8%) | 0.973 |
| Married | 25 (38.5%) | 47 (38.2%) |  |
| **Pre-injury residence\*\*** |  |  |  |
| Home (live alone) | 14 (21.2%) | 31 (24.8%) | 0.710 |
| Home (live with family) | 50 (75.8%) | 92 (73.6%) |  |
| Institution | 2 (3.0%) | 2 (1.6%) |  |
| **BMI\*** | 23.82 ± 4.77 | 25.60 ± 3.91 | 0.008 |
| **CCI groups\*\*** |  |  |  |
| No comorbidity/mild | 21 (31.8%) | 68 (54.4%) | 0.009 |
| Moderate | 36 (54.6%) | 42 (33.6%) |  |
| Severe | 9 (13.6%) | 15 (12.0%) |  |
| **SPMSQ\*** | 6.79 ± 1.67 | 7.83 ± 1.63 | <0.001 |
| **EQ5D before fracture\*** | 0.73 ± 0.17 | 0.83 ± 0.14 | <0.001 |
| **Barthel index before fracture\*** | 92.95 ± 7.70 | 96.16 ± 5.54 | 0.003 |
| **Type of fracture\*\*** |  |  |  |
| Femoral neck | 25 (37.9%) | 59 (47.2%) | 0.462 |
| Intertrochanteric | 35 (53.0%) | 57 (45.6%) |  |
| Subtrochanteric | 6 (9.1%) | 9 (7.2%) |  |
| **Time from admission to operation\*** | 6.26 ± 3.17 | 5.75 ± 2.98 | 0.277 |
| **Lenght of hospital stay\*** | 15.91 ± 5.20 | 16.03 ± 4.42 | 0.864 |
| **Surgical procedure\*\*** |  |  |  |
| Arthroplasty | 27 (36.4%) | 57 (45.6%) | 0.219 |
| ORIF | 39 (63.6%) | 68 (54.4%) |  |
| **Type of anaesthesia\*\*** |  |  |  |
| General | 51 (79.7%) | 78 (64.5%) | 0.032 |
| Regional | 13 (20.3%) | 43 (35.5%) |  |
| **Duration of anaesthesia\*** | 118.47 ± 39.54 | 114.70 ± 27.18 | 0.504 |
| **Complications\*\*** |  |  |  |
| Yes | 18 (27.3%) | 30 (24.0%) | 0.620 |
| No | 48 (72.7%) | 95 (76.0%) |  |

\*Values are given as the mean with the standard deviation in parentheses
\*\* Values are given as the number of patients with the percentage in parentheses
RR - relative risk; BMI - body mass index; CCI - Charlson Comorbidity Index; SPMSQ - Short Portable Mental Status
Questionnaire; HGS - Handgrip strength

Patients with relevant hand grip weakness achieved statistically significant lower Barthel index scores 3 (56.30 ±25.87 vs. 75.77 ±21.49) (Table 2) and 6 months (67.77 ±29.15 vs. 87.66 ±19.30) (Table 3) after hip fracture. Adjusted multivariate regression analysis revealed that hand grip strength below the cutoff point for sarcopenia according to the EWGSOP2 was an independent predictor of worse functional outcome at 3 and 6 months after hip fracture for both genders and in all age populations.

**TABLE 2.** Univariate and Multivariate analysis for variables significantly associated with Barthel index scores 3 months after fracture

| Predictors | Univariate analysis |  | Multivariate analysis |  |
| --- | --- | --- | --- | --- |
|  | B (95% CI) | p value | B (95% CI) | p value |
| **Marital status** | -0.05 (-11.04 – 5.71) | 0.531 |  |  |
| **Preinjury residence** | -0.18 (-17.22 - -1.41) | 0.021 | -0.13 (-13.29 - -0.34) | 0.039 |
| **BMI** | 0.02 (-0.79 – 1.08) | 0.761 |  |  |
| **CCI** | -0.29 (-15.40 - -5.24) | <0.001 | -0.10 (-8.12 – 1.06) | 0.131 |
| **SPMSQ** | 0.27 (1.72 – 5.97) | <0.001 | 0.72 (-0.92 – 2.94) | 0.302 |
| **EQ5D before fracture** | 0.31 (25.94 – 71.55) | <0.001 | 0.16 (2.10 – 46.10) | 0.032 |
| **Barthel index before fracture** | 0.51 (1.37 – 2.30) | <0.001 | 0.36 (0.77 – 1.71) | <0.001 |
| **HGS** | 0.31 (8.29 – 24.51) | <0.001 | 0.185 (2.29 – 16.71) | 0.010 |
| **Time from admission to operation** | -0.01 (-1.36 – 1.13) | 0.860 |  |  |
| **Lenght of hospital stay** | -0.04 (-1.04 – 0.62) | 0.616 |  |  |
| **Type of anaesthesia** | 0.09 (-3.22 – 12.55) | 0.244 |  |  |
| **Duration of anaesthesia** | -0.21 (-0.31 – 0.05) | 0.009 | -0.12 (-0.20 – 0.01) | 0.069 |
| **Complications** | -0.16 (-18.58 - -0.44) | 0.040 | -0.13 (-14.90 - -0.20) | 0.044 |

Adjusted for age and gender
BMI - body mass index; CCI - Charlson Comorbidity Index; SPMSQ - Short Portable Mental Status Questionnaire; HGS - Handgrip strength

**TABLE 3.** Univariate and Multivariate analysis for variables significantly associated with Barthel index scores 6 months after fracture

| **Predictors** | **Univariate analysis** |  | **Multivariate analysis** |  |
| --- | --- | --- | --- | --- |
|  | B (95% CI) | p value | B RR (95% CI) | p value |
| **Marital status** | -0.12 (-14.75 – 2.56) | 0.166 |  |  |
| **Preinjury residence** | -0.14 (-15.52 – 1.20) | 0.093 |  |  |
| **BMI** | 0.04 (-0.75 – 1.28) | 0.602 |  |  |
| **CCI** | -0.31 (-16.61 - -5.58) | <0.001 | -0.17 (-10.99 - -1.03) | 0.018 |
| **SPMSQ** | 0.29 (1.84 – 6.30) | <0.001 | 0.09 (-0.76 – 3.26) | 0.222 |
| **EQ5D before fracture** | 0.36 (32.11 – 78.99) | <0.001 | 0.15 (-0.28 – 45.56) | 0.053 |
| **Barthel index before fracture** | 0.53 (1.47 – 2.48) | <0.001 | 0.38 (0.86 – 1.91) | <0.001 |
| **HGS** | 0.36 (10.39 – 27.18) | <0.001 | 0.21 (3.07 – 18.37) | 0.006 |
| **Time from admission to operation** | -0.07 (-1.87 – 0.74) | 0.392 |  |  |
| **Lenght of hospital stay** | 0.01 (-0.83 – 0.94) | 0.895 |  |  |
| **Type of anaesthesia** | 0.08 (-4.19 – 12.28) | 0.333 |  |  |
| **Duration of anaesthesia** | -0.17 (-0.28 - -0.01) | 0.037 | -0.08 (-0.17 – 0.05) | 0.250 |
| **Complications** | -0.07 (-14.06 – 5.82) | 0.414 |  |  |

Adjusted for age and gender
BMI - body mass index; CCI - Charlson Comorbidity Index; SPMSQ - Short Portable Mental Status Questionnaire; HGS -
Handgrip strength

Besides hand grip strength. living at home, better quality of life and higher functionally independence before fracture, as well as absence of complications during the hospitalization period were independent predictors of Barthel index scores 3 months postoperatively (Table 2). Multivariate regression analysis showed that, besides hand grip strength above cutoff values for sarcopenia, lower CCI index and higher Barthel index scores before fracture were independent predictors of higher Barthel index scores 6 months after fracture (Table 3).

## Discussion

Our study demonstrates that hip fracture patients with a validated threshold for clinical weak grip strength assessed at an early stage had significantly poorer functional recovery after 3 and 6 months compared to patients with a grip strength above the cutoff points. Furthermore, our findings provide evidence that hand grip strength along with several other prognostic factors traditionally considered in clinical practice can independently predict short- and long-term functional outcome [25]. Within the study 35% of the study population had relevant clinical weakness based on hand grip strength. Older age, higher level of comorbidity, lower cognitive level, lower functional level, and worse quality of life recognized at admission indicate clearly a decline of reserve and function across multiple physiological systems in this group of patients.

The results of our investigation are consistent with data from previous studies confirming the prognostic role of handgrip strength [12–17]. However, it is not easy to compare present results with other studies addressing this subject. First, handgrip strength was assessed at various time points in different studies. There are only two studies evaluating the prognostic value of handgrip strength measured in the acute setting [16, 17]. Savino et al. showed that handgrip strength measured at hospital admission significantly predicted walking recovery 12 months after hip [16]. Alvarez MN et al. concluded that HGS asessed in the first hours after hospital admission for hip fracture surgery is an indicator of functional recovery after three months [17]. It is well known that muscle mass is maintained during the first 10 days, although it subsequently diminishes [26, 27]. Consequently, it is reasonable to assume that measuring HGS early after hip fracture is an appropriate time to assess function. Second, a small body of literature used cutoff points to define clinical relevant weakness based on HGS [15, 17, 28], and no study applied the EWGSOP2 criteria. Alvarez MN et al. [17] who used the EWGSOP criteria [11], and di Monaco et al. [15] who used the FNIH Sarcopenia Project criteria for HGS categorization [29] confirmed the prognostic role of hand grip strength. In contrast, Steihaug et al. who applied the EWGSOP criteria to investigate the impact of HGS early after fracture were the only ones who found no association between grip strength and short- and long-term functional outcome [28]. This is the only study to deny the value of HGS in predicting functional outocome in hip fracture patients. It has to be taken into account that although the definition of „weakness“ according to the FNIH, and the EWGSOP criteria correspond very closely to the one defined by the EWGSOP2 [3], it is not the same. Moreover, our results cannot be compared completely to those published by de Monaco et al., because they reported their results only on women, and in the postacute rehabilitation setting [13, 15].

There are several strengths of our study. First, to the best of our knowledge this is the first study to assess HGS using the EWGSOP2 criteria. Second, our study proves the prognostic value of HGS in the acute setting for both gender and all ages. Most studies who analyzed the predictive role of hand grip strength reported their results only in women with hip fracture [12, 13, 15].

There are also some limitations to our study. First, the outcome of our study was assessed with only self-reported information collected by phone interviews. Second, patients were collected only from one single center. Additionally, there are other confounding factors that could have been studied, for example nutritional status and vitamin D status.

Our results have two clinical implications. First, assessment of HGS can be used to identify hip fracture patients at high risk for poor functional outcome at an early time point. Second, it is well known that functional evaluation is hard to assess in acute hip fracture patients. In the first place, gait speed cannot be assessed before surgery. Thus, functional evaluation is limited to measuring muscle strength. Confirmation of the prognostic value of HGS assessed in the acute setting is therefore very significant. Third, muscle weakness is a modifiable risk factor that can be improved. It is well know that strengthening exercises had favorable effects on various outcomes after hip fracture [30]. Therefore, future studies should reveal if patients with clinically defined weakness who sustain a hip fracture could benefit from interventions to improve muscle strength, function, and outcomes.

Our study has identified HGS assessed in the acute setting as potential prognostic predictor of functional outcome in patients with hip fracture. Hand grip strength is an accessible, cost effective, and simple objective measure of physical function for bedridden patients. Thus, clinicians should be encouraged to include hand grip assessment in their evaluation of hip fracture patients at admission to the acute setting in order to optimize prognostic counseling and treatment of high-risk individuals. Further studies are needed to investigate the relevance of early introduction of resisting exercise programs in patients hip fracture patients with relevant low muscle strength.

## Acknowledgments

This study is conducted as the part of project No 175046 funded by Ministry of Education, Science and Technological Development, Republic of Serbia, 2011-2020.

## References

1. Bandeen-Roche K, Xue QL, Ferrucci L, Walston J, Guralnik JM, Chaves P, et al. Phenotype of frailty: characterization in the women’s health and aging studies. J Gerontol A Biol Sci Med Sci. 2006 Mar;61(3):262–6.

2. Fielding RA, Vellas B, Evans WJ, Bhasin S, Morley JE, Newman AB, et al. Sarcopenia: an undiagnosed condition in older adults. Current consensus definition: prevalence, etiology, and consequences. International working group on sarcopenia. J Am Med Dir Assoc. 2011 May;12(4):249–56.

3. Cruz-Jentoft AJ, Bahat G, Bauer J, Boirie Y, Bruyère O, Cederholm T, et al. Sarcopenia: revised European consensus on definition and diagnosis. Age Ageing. 2019 Jan 1;48(1):16–31.

4. Landi F, Calvani R, Ortolani E, Salini S, Martone AM, Santoro L, et al. The association between sarcopenia and functional outcomes among older patients with hip fracture undergoing in-hospital rehabilitation. Osteoporos Int. 2017 May;28(5):1569–1576.

5. He H, Liu Y, Tian Q, Papasian CJ, Hu T, Deng HW. Relationship of sarcopenia and body composition with osteoporosis. Osteoporos Int. 2016 Feb;27(2):473–82.

6. Benichou O, Lord SR. Rationale for Strengthening Muscle to Prevent Falls and Fractures: A Review of the Evidence. Calcif Tissue Int. 2016 Jun;98(6):531–45.

7. Hirschfeld HP, Kinsella R, Duque G. Osteosarcopenia: where bone, muscle, and fat collide. Osteoporos Int. 2017 Oct;28(10):2781–2790.

8. Rantanen T, Volpato S, Ferrucci L, Heikkinen E, Fried LP, Guralnik JM. Handgrip strength and cause-specific and total mortality in older disabled women: exploring the mechanism. J Am Geriatr Soc. 2003 May;51(5):636–41.

9. Syddall H, Cooper C, Martin F, Briggs R, Aihie Sayer A. Is grip strength a useful single marker of frailty? Age Ageing. 2003 Nov;32(6):650–6.

10. Chen LK, Liu LK, Woo J, Assantachai P, Auyeung TW, Bahyah KS, et al. Sarcopenia in Asia: consensus report of the Asian Working Group for Sarcopenia. J Am Med Dir Assoc. 2014 Feb;15(2):95–101.

11. Cruz-Jentoft AJ, Baeyens JP, Bauer JM, Boirie Y, Cederholm T, Landi F, et al. Sarcopenia: European consensus on definition and diagnosis: Report of the European Working Group on Sarcopenia in Older People. Age Ageing. 2010 Jul;39(4):412–23.

12. Wehren LE, Hawkes WG, Hebel JR, Orwig DL, Magaziner J. Bone mineral density, soft tissue body composition, strength, and functioning after hip fracture. J Gerontol A Biol Sci Med Sci. 2005 Jan;60(1):80–4.

13. Di Monaco M, Castiglioni C, De Toma E, Gardin L, Giordano S, Di Monaco R, et al. Handgrip strength but not appendicular lean mass is an independent predictor of functional outcome in hip-fracture women: a short-term prospective study. Arch Phys Med Rehabil. 2014 Sep;95(9):1719–24.

14. Beloosesky Y, Weiss A, Manasian M, Salai M. Handgrip strength of the elderly after hip fracture repair correlates with functional outcome. Disabil Rehabil. 2010;32(5):367–73.

15. Di Monaco M, Castiglioni C. Weakness and Low Lean Mass in Women With Hip Fracture: Prevalence According to the FNIH Criteria and Association With the Short-Term Functional Recovery. J Geriatr Phys Ther. 2017 Apr/Jun;40(2):80–85.

16. Savino E, Martini E, Lauretani F, Pioli G, Zagatti AM, Frondini C, et al. Handgrip strength predicts persistent walking recovery after hip fracture surgery. Am J Med. 2013 Dec;126(12):1068–75.e1.

17. Alvarez MN, Bonnardeaux P L.D, Thuissard IJ, Sanz-Rosa D, Muñana EA, Galindo RB, et al. Grip strength and functional recovery after hip fracture: An observational study in elderly population. Eur Geriatr Med. 2016 Dec 1;7(6):556–60.

18. Pfeiffer E. A short portable mental status questionnaire for the assessment of organic brain deficit in elderly patients. J Am Geriatr Soc. 1975 Oct;23(10):433–41.

19. Richards L G. Posture effects on grip strength. Arch Phys Med Rehabil. 1997 Oct;78(10):1154–6.

20. Katz S, Ford AB, Moskowitz RW, Jackson BA, Jaffe MW. Studies of Illness in the Aged. The Index of ADL: A Standardized Measure of Biological and Psychological Function. JAMA. 1963 Sep 21;185:914–9.

21. Mahoney FI, Barthel D. Functional evaluation: The Barthel Index. Md State Med J. 1965 Feb;14:61–5.

22. Brooks R. EuroQol: the current state of play. Health Policy. 1996 Jul;37(1):53–72.

23. Dolan P. Modeling valuations for EuroQol health states. Med Care. 1997 Nov;35(11):1095–108.

24. Charlson ME, Pompei P, Ales KL, MacKenzie CR. A new method of classifying prognostic comorbidity in longitudinal studies: development and validation. J Chronic Dis. 1987;40(5):373–83.

25. Kristensen MT. Factors affecting functional prognosis of patients with hip fracture. Eur J Phys Rehabil Med. 2011 Jun;47(2):257–64.

26. D’Adamo CR, Hawkes WG, Miller RR, Jones M, Hochberg M, Yu-Yahiro J, et al. Short-term changes in body composition after surgical repair of hip fracture. Age Ageing. 2014 Mar;43(2):275–80.

27. Fox KM, Magaziner J, Hawkes WG, Yu-Yahiro J, Hebel JR, Zimmerman SI, et al. Loss of bone density and lean body mass after hip fracture. Osteoporos Int. 2000;11(1):31–5.

28. Steihaug OM, Gjesdal CG, Bogen B, Kristoffersen MH, Lien G, Ranhoff AH. Sarcopenia in patients with hip fracture: A multicenter cross-sectional study. PLoS One. 2017 Sep 13;12(9):e0184780.

29. Alley DE, Shardell MD, Peters KW, McLean RR, Dam TT, Kenny AM, et al. Grip strength cutpoints for the identification of clinically relevant weakness. J Gerontol A Biol Sci Med Sci. 2014 May;69(5):559–66.

30. Lee SY, Yoon BH, Beom J, Ha YC, Lim JY. Effect of Lower-Limb Progressive Resistance Exercise After Hip Fracture Surgery: A Systematic Review and Meta-Analysis of Randomized Controlled Studies. J Am Med Dir Assoc. 2017 Dec 1;18(12):1096.e19-1096.e26.

